# Study of holobiont resilience following a break in symbiotic transmission in two rabbit lines selected for feed efficiency

**DOI:** 10.64898/2026.09.04.749168

**Authors:** Hervé Garreau, Marie Renevey, Patrick Aymard, Laurent Cauquil, Virginie Helies, Catherine Larzul, Maryse Poli, Julien Ruesche, Olivier Zemb, Sylvie Combes

## Abstract

This study was carried out on two rabbit lines, each selected for a feed-efficiency criterion (growth on restricted feed for the AlimR line or residual consumption on an ad libitum diet for the ConsR line), introduced into a facility by adoption of newborn kits by SPF (specific-pathogen-free) females. Following this break in symbiotic transmission, fecal samples from females were used to study the microbiota composition of the two lines over seven generations (17 females per line and per generation). The proportion of Firmicutes was significantly higher in the AlimR line than in the ConsR line (82.22 % vs 78.21 %, p < 0.0001). Conversely, the proportion of Bacteroidota was significantly higher in the ConsR line than in the AlimR line (20.38 % vs 16.50 %, p < 0.0001). The effect of generation was also significant for the abundance of Firmicutes and Bacteroidota (p = 0.0009 and p < 0.0001, respectively). The Shannon diversity index was significantly higher in the AlimR line than in the ConsR line (5.23 vs 5.14, p = 0.02). The effect of generation was also significant for the Shannon index (p = 0.003). Finally, a discriminant principal component analysis of the microbiota composition allowed us to distinguish both the two lines and the generations, in particular the 11th selected generation (animals adopted by SPF females) and the 18th generation.

## Introduction

The rabbit digestive tract harbors a complex and diverse microbial community that plays a key role in many physiological functions such as digestion and immunity (Combes et al. [2]). Manipulating the microbiota appears to be a promising avenue for improving the sustainability of rabbit production, especially in terms of reducing pharmaceutical inputs and enhancing feed efficiency. Studies carried out in other species have demonstrated the influence of numerous factors on gut microbiota composition: rearing environment (diet composition and amount, humidity, temperature), maternal transmission, age and physiological state, genetics, and antibiotic treatments (Gilbert et al. [6]). In rabbits, maternal transmission of the microbiota to the young is well documented, with ingestion of maternal caecotrophes in the nest having been demonstrated (Combes et al. [3]). Genetic control of microbiota composition has been highlighted by differences observed between divergent lines selected for feed efficiency in pigs (Aliakbari et al. [1]) or for digestive efficiency in chickens (Mignon-Grasteau et al. [10]).

To better understand holobiont assembly and host–microbiota interaction mechanisms, it is essential to set up experimental designs that avoid confounding genetic and environmental factors by controlling each component. Two experimental rabbit lines, each selected for a feed-efficiency criterion, were introduced into a renovated building of the INRAE experimental unit (UMR GenPhySE) by adopting newborn kits from Specific Pathogen Free (SPF) females. Starting from a break in symbiotic transmission and using a common rearing environment for the two lines, the aim of this experiment was to study, using fecal samples from breeding females, the host–microbiota interactions (genotype effect on the microbiota) and the holobiont’s resilience across generations.

## Materials and Methods

### Animals

The animals originated from the INRA 1001 line (Larzul & De Rochambeau, 2005) and were raised at the INRAE experimental facility of UMR GenPhySE (Castanet-Tolosan, France) in compliance with the French animal experimentation regulations (APAFiS licence #18416). Two lines were used in this study:

1. AlimR – selected for rapid growth under restricted feeding (80 % of ad libitum level).
2. ConsR – selected for low residual consumption, i.e. total feed intake under ad libitum conditions corrected (by linear regression) for average metabolic body weight and for daily weight gain between 30 and 63 days.

Both lines are reared simultaneously in the same maternal and growth cages. For each generation, females of the two lines are inseminated four times with a 42-day interval. At each generation, 300 animals originating from inseminations 3 and 4 are monitored individually in cages from weaning (30 days) to 63 days of age. For both lines, body weight measured at weaning (30 days) and at 63 days allowed the calculation of the average daily gain (ADG). Individual feed intakes from 30 to 63 days were used to compute residual consumption. Animals monitored during growth are selected according to their estimated breeding value (EBV) for ADG (AlimR) or for residual consumption (ConsR) obtained with the ASReml software (Gilmour et al. [7]). The detailed protocol is described by Drouilhet et al. [4]. From the 1st to the 10th generation, each line comprised 54 females and 27 males (one sire and two replacements), distributed in nine breeding groups.

### Break in Symbiotic Transmission

To renovate the selection building, the 9th generation of each line was transferred to an annex building. Generations 9 and 10 were raised in this annex following the same management and selection procedures as in the main facility. To improve the sanitary status of the lines, a herd of 150 pregnant Specific Pathogen Free females from Charles River Laboratories was installed in the renovated selection building to receive the kits of the 11th generation. Kits resulting from the 5th artificial insemination of the 10th-generation females were dusted with antibiotic-impregnated talc and, immediately after birth, transferred from the annex to the renovated selection building:

1. AlimR – 62 male and 96 female kits were adopted by 31 EOPS females.
2. ConsR – 93 male and 119 female kits were adopted by 39 EOPS females.

After individual performance assessment and calculation of selection index, 84 females and 27 males per line were retained to produce the 12th generation. Generations 12 through 18 were carried out and selected using the same protocol described in section 2.1.

### Determination of Microbiota Composition

In each line and for each generation from the 11th to the 18th, feces from 17 females (1–2 per family) were collected two days before the third artificial insemination and stored at –80 °C. Due to a storage issue, samples from generation 17 were excluded from the analysis. In total, 232 samples were processed. Microbial DNA was extracted with the ZR Soil Microbe DNA MiniPrep™ kit (ZymoResearch, Freiburg, Germany). The bacterial 16S rRNA gene V3-V4 hypervariable regions were amplified and sequenced using an Illumina MiSeq platform (Get-PlaGe, Toulouse Occitanie, doi:10.15454/1.5572370921303193E12). V3-V4 regions were chosen because they provide the most precise taxonomic assignment (Yang et al. [14]). Sequences were processed with the FROGS pipeline (Escudié et al. [5]), which employs the Swarm algorithm for clustering (Mahé et al. [9]). Chimeric sequences were removed with VSEARCH (Rognes et al. [12]). Clusters representing less than 0.005 % of total reads (4 288 674 reads) were discarded. Taxonomic assignment was performed with BLAST against the SILVA SSU Ref NR 132 database (Quast et al. [11]). An R phyloseq object (McMurdie & Holmes, 2013) was generated and alpha-diversity indices were calculated after rarefying to 5 980 reads (observed OTU count and Shannon index).

### Statistical Analyses

A variance analysis was performed to compare the proportion of each phylum and the alpha-diversity indices using R 4.1.2. Fixed effects in the model were line (n = 2) and generation (n = 7). The effects of line and generation on microbiota composition were explored with a discriminant analysis of principal components (DAPC) (Jombart et al. [8]).

## Results and Discussion

### Taxonomic Composition of the Microbiota

The proportion of Firmicutes, the most abundant phylum in rabbits, was significantly higher in the AlimR line than in the ConsR line (82.22 % vs 78.21 %, *p* < 0.0001; Figure 1). Conversely, the proportion of Bacteroidota, the second most abundant phylum, was significantly higher in the ConsR line (20.38 % vs 16.50 %, *p* < 0.0001). No line effect was observed for the other phyla (Actinobacteriota, Desulfobacterota, Proteobacteria) whose relative abundances were respectively 0.66 %, 0.11 %, 0.60 % in AlimR and 0.66 %, 0.12 %, 0.73 % in ConsR. The generation effect was significant only for Firmicutes and Bacteroidota (*p* = 0.0009 and *p* < 0.0001, respectively). From generation 11 to 18, the proportion of Firmicutes generally increased (from 79.7 % to 86.3 % in AlimR and from 80.0 % to 83.2 % in ConsR), whereas Bacteroidota decreased (from 19.0 % to 11.4 % in AlimR and from 18.9 % to 14.6 % in ConsR). The line × generation interaction was not significant. Independently of line or generation, the predominance of these two phyla in rabbit gut microbiota has been previously reported (Combes et al. [2] ; Velasco-Galilea et al., 2018).

**Figure 1:**
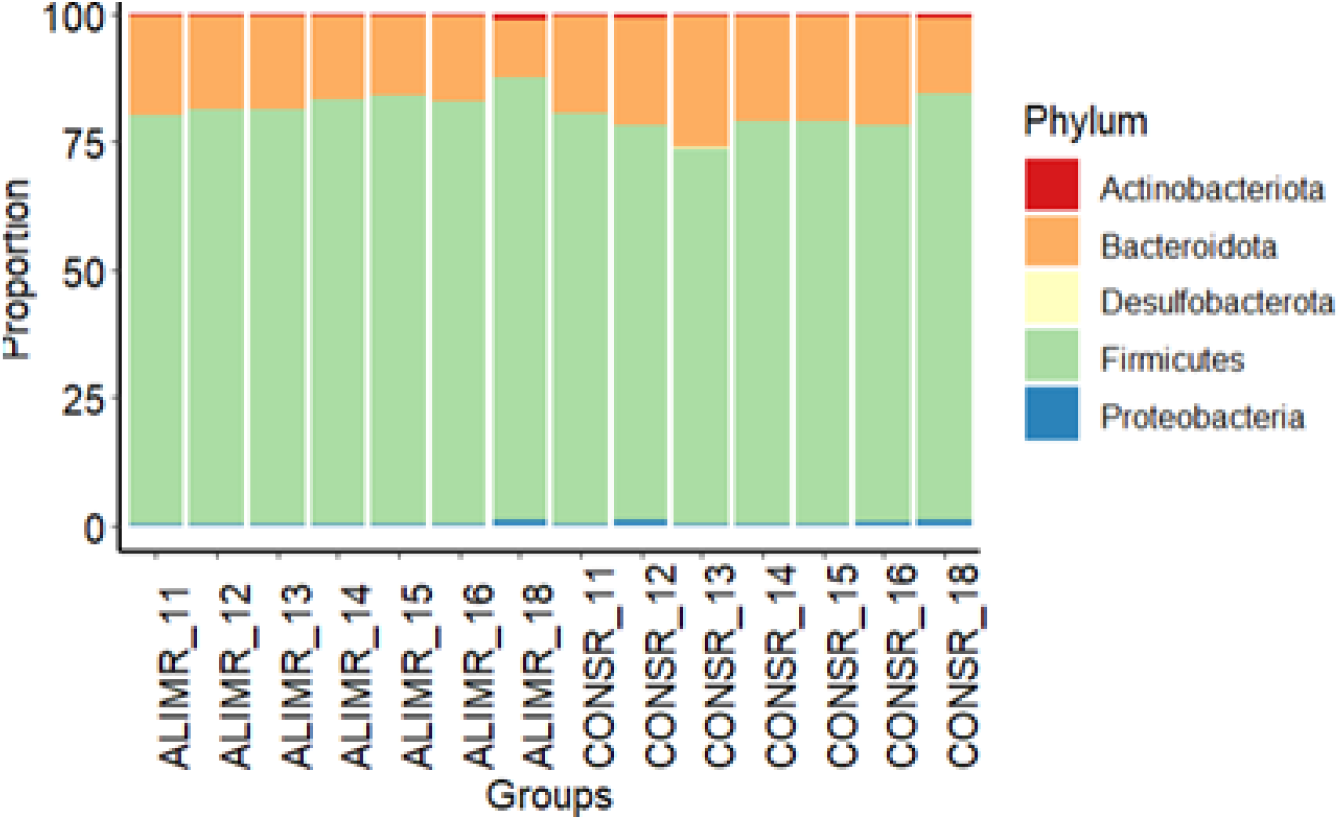
Proportion of each phylum for the AlimR and ConsR lines across generations 11 to 18.

### Microbiota Diversity

Alpha-diversity indices by line and generation are shown in Figure 2. The observed OTU count did not differ significantly between lines (641 in AlimR vs 631 in ConsR). However, the Shannon index, which weighs proportional species abundance, was significantly higher in the AlimR line (selected for growth under restriction) than in the ConsR line (selected for low residual consumption) (5.23 vs 5.14, *p* = 0.02). In pigs, a study comparing microbial diversity of two divergent lines selected for residual feed intake found the opposite pattern, with higher Shannon diversity in the low-consumption line (Aliakbari et al., 2021). The generation effect was also significant for the Shannon index (*p* = 0.003): generation 18 showed the highest values (5.40 in AlimR and 5.35 in ConsR), whereas generations 13 and 14 displayed the lowest values (5.21 and 5.19, respectively, in AlimR and ConsR).

**Figure 2:**
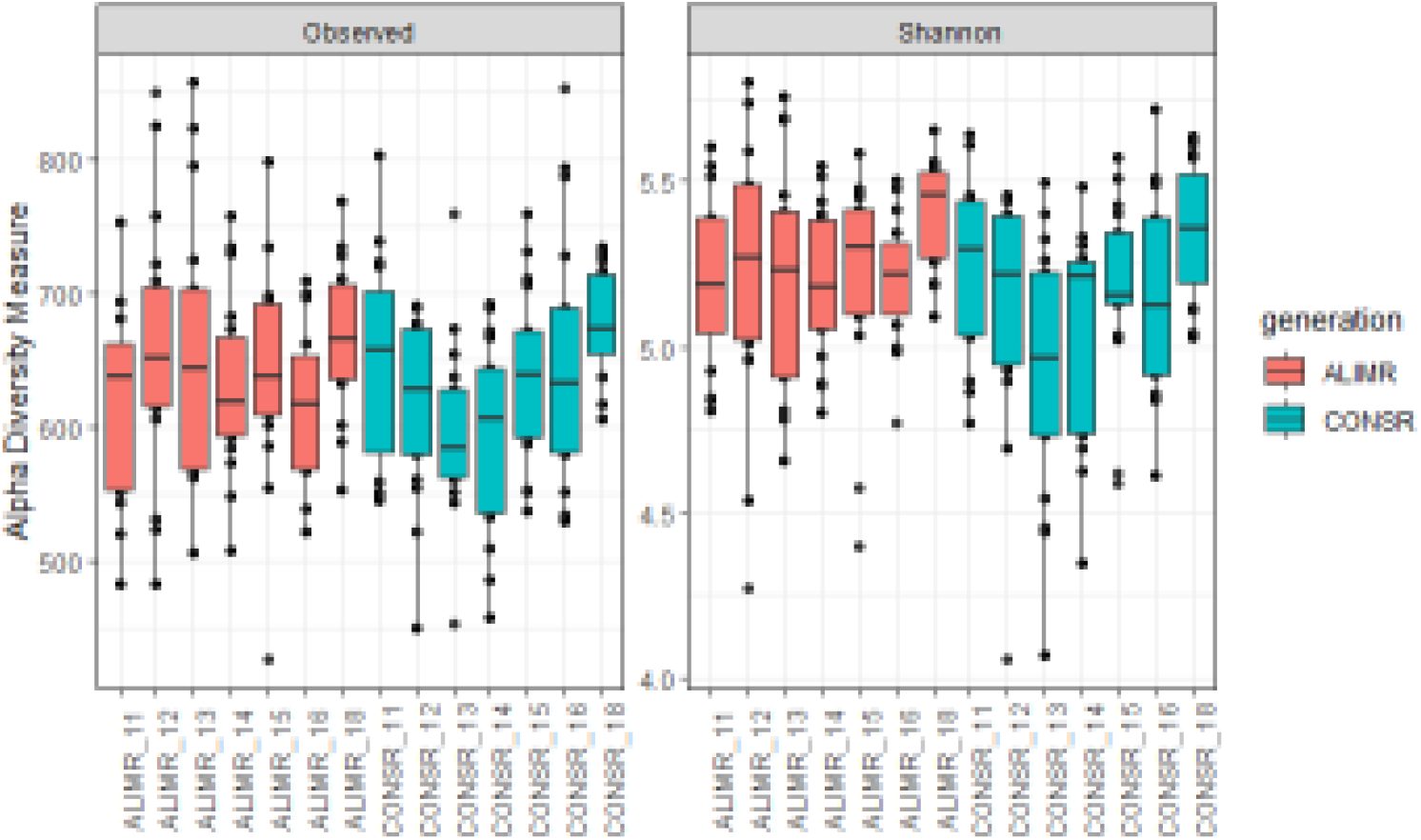
Observed richness and Shannon index for the AlimR and ConsR lines across generations 11 to18.

### Discriminant Principal Component Analysis of Microbiota Composition

DAPC of the microbiota composition clearly separated the two lines (Figure 3). Similar line-related microbiota differences have been reported in pigs selected for feed efficiency (Aliakbari et al., 2021) and in chickens selected for digestive efficiency (Mignon-Grasteau et al. [10]). The persistence of line-specific microbiota signatures beyond the break in maternal transmission underscores a direct effect of host genetics on microbiota control. Human metagenome-wide association studies (mGWAS) on large cohorts have identified preferential associations between a limited set of taxa and single-nucleotide polymorphisms (Turpin et al. [13]). Nevertheless, the physiological mechanisms governing host-driven selection remain to be elucidated.

**Figure 3:**
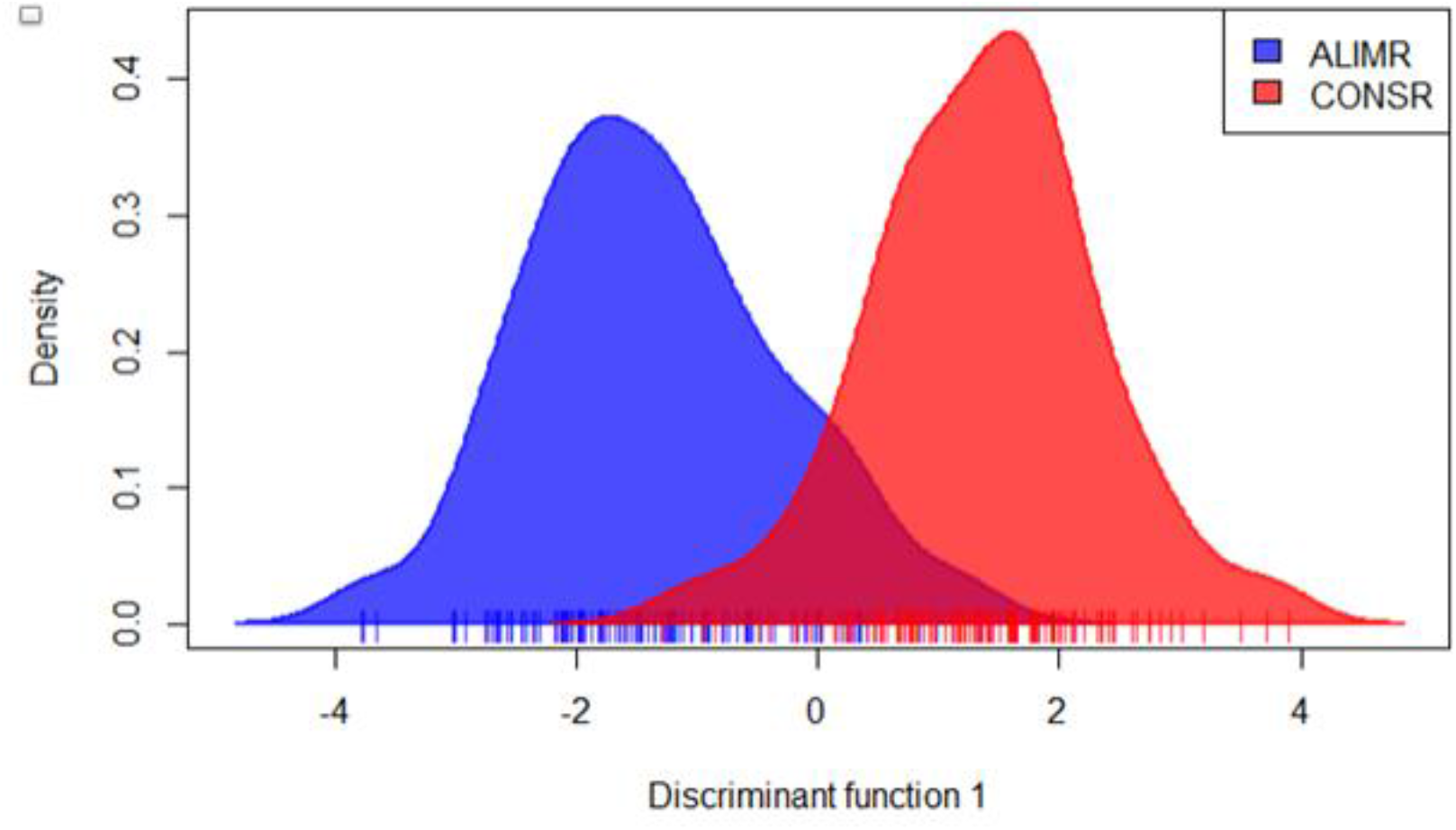
Kernel density estimation of DAPC for the AlimR and ConsR microbiota data.

When DAPC was performed with generation as the grouping factor (Figure 4), the 11th generation—corresponding to kits adopted by SPF females—differed from generations 12 to 16, which displayed similar microbial profiles. The 18th generation also formed a distinct cluster. No change in animal management that could explain the particular microbiota profile of generation 18 was identified.

**Figure 4:**
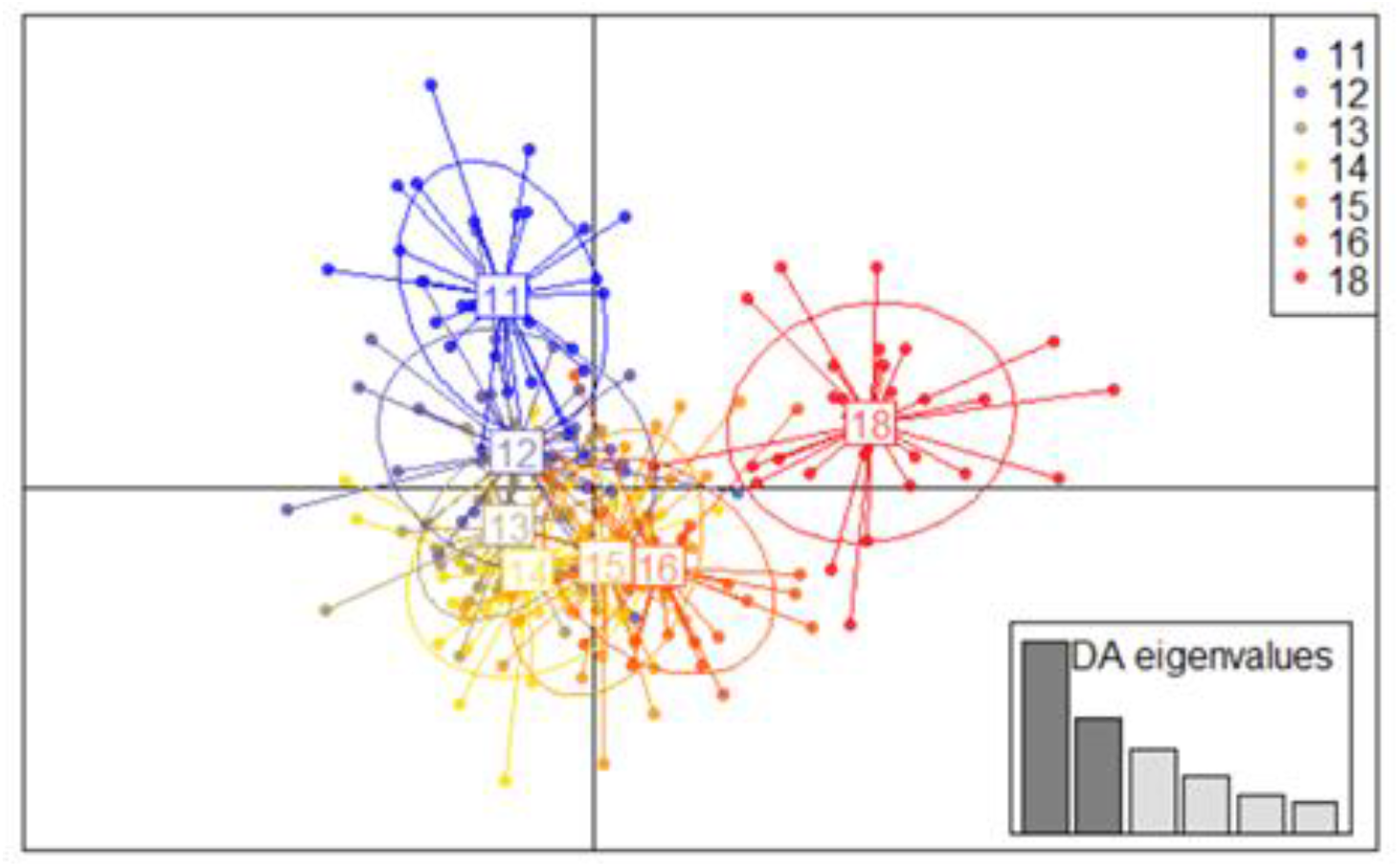
DAPC of microbiota data for each generation across all animals.

## Conclusions

Following a break in symbiotic transmission caused by adoption of kits by SPF (EOPS) females, we demonstrated clear differences in taxonomic profiles, diversity, and fecal microbiota composition between two rabbit lines selected for opposite feed-efficiency criteria. Generation effects were also significant for phylum proportions, diversity indices, and overall microbiota composition, notably highlighting marked differences for the 11th generation (adopted by SPF females) and the 18th generation.

## Acknowledgments

We thank the staff of the GenPhySE experimental facility for their care in animal husbandry and data recording, the Genotoul bioinformatics platform (Bioinfo Genotoul, doi : 10.15454/), and the SIGENAe group for computational and storage support.

## Notes

### Competing Interest Statement

The authors have declared no competing interest.

